# *APOE* genotype or presence of brain amyloid alters the plasma proteome in cognitively normal, elderly subjects

**DOI:** 10.1101/2022.12.28.522119

**Authors:** Sarah M. Philippi, BP Kailash, Towfique Raj, Joseph M. Castellano

**Affiliations:** Nash Family Department of Neuroscience, Department of Neurology, Friedman Brain Institute, Icahn School of Medicine at Mount Sinai, New York, NY, United States; Ronald M. Loeb Center for Alzheimer’s Disease, Icahn School of Medicine at Mount Sinai, New York, NY, United States; Graduate School of Biomedical Sciences, Icahn School of Medicine at Mount Sinai, New York, NY, United States; Black Family Stem Cell Institute, Icahn School of Medicine at Mount Sinai, New York, NY, United States; Department of Genetics and Genomic Sciences, Icahn Institute for Data Science and Genomic Technology, Icahn School of Medicine at Mount Sinai, New York, NY, United States

**Keywords:** APOE, Alzheimer’s disease, plasma proteins, rejuvenation, plasma proteome, amyloid, SOMAscan, extracellular matrix, inflammation

## Abstract

**Background:** Processes that drive Alzheimer’s disease pathogenesis have long been considered to occur within the central nervous system, yet recent studies have bolstered the possibility that changes in the periphery may be relevant to the disease process. Accumulating evidence has suggested that proteins changing in the blood may be reliable indicators of disease within the brain. Recent advances in geroscience have identified potential mechanisms of blood-brain communication that modulate brain function in ways that could be harnessed for therapy. While blood-borne proteins associated with either youth or old age have been targeted to restore function to the aged brain, it remains unclear whether other dysfunctional systemic states can be exploited for similar benefits. Here we investigate whether *APOE* allelic variation or presence of brain amyloid are associated with distinct proteomic changes within the systemic environment and what molecular processes are associated with these changes.

**Methods:** Using the SOMAscan assay, we measured 1,305 plasma proteins from 53 homozygous *APOE3* and *APOE4* subjects (mean age = 68 years; minimum = 54 years) who exhibited no cognitive impairment, some of whom can be categorized as harboring cerebral amyloid based on cerebrospinal fluid Aβ42 measurements. Using the Dream R package for linear mixed effects modeling, we investigated possible contributions of either the *APOE-ε4* allele or amyloid positivity to changes in the plasma proteome. Ontology-based pathway and module trait correlation analyses were performed to understand disrupted pathways that vary based on *APOE* genotype or amyloid positivity.

**Results:** We found that expression of the *APOE-ε4* allele produced distinct changes in the composition of the plasma proteome. Using both pathway enrichment analysis and weighted gene co-expression network analysis, we found that plasma proteins associated with *APOE4* expression were linked to pathways related to atherosclerosis, lipid transport, the extracellular matrix, and synaptogenesis signaling. Independent of *APOE4*, we found that cognitively normal, amyloid-positive subjects exhibit distinct plasma proteome signatures associated with pathways previously linked to AD pathology, relative to amyloid-negative controls. Harboring brain amyloid was associated with plasma proteomic changes linked to dysfunction in blood-brain barrier and other neural cell types. Our results indicate that changes in the plasma proteome are related to possession of AD risk alleles, as well as the presence of amyloid pathology in subjects prior to the onset of symptoms. This work highlights the possibility that pathways in the systemic environment in certain risk contexts may be plausible targets to explore for modulating disease.

## Background

Alzheimer’s disease (AD) is a devastating neurodegenerative disease for which aging serves as the strongest risk factor. The development of AD is associated with several hallmark pathologies, including the accumulation of amyloid-β into plaques, which can be detected in living subjects using a variety of reliable biomarkers (1-5). To date, there are no effective treatments to substantially change the course of disease despite the increasing toll it will have as the global population ages. One intriguing direction has been to explore how aging acts as a risk factor for AD, particularly through characterization of blood-CNS communication pathways that have been shown to be perturbed in mice and humans (6-12). These studies support the idea that age-associated changes in the plasma proteome can be harnessed to revitalize age-sensitive organ systems (7, 9, 10, 12-17). Our group and others have demonstrated the sufficiency of youth-associated blood-borne proteins, via peritoneal parabiosis or through plasma transfer, to revitalize the aged mouse brain and restore hippocampal function (7). Intriguingly, we found that human plasma proteins were sufficient to recapitulate these phenotypes in aged mice (7). While these studies demonstrate that brain function can be modified by aging-associated blood-borne proteins, it remains unclear whether pathological processes within the AD brain alter the systemic environment or whether risk factors modulate the composition of the blood proteome that may ultimately affect brain health.

The strongest genetic risk factor for sporadic, late-onset AD is possession of the *APOE-ε4* allele (18, 19). Compared to the common *APOE-ε3* allele, the *APOE-ε4* allele markedly increases AD risk 3-12 fold (18-20), depending on the number of copies expressed. It is well established that *APOE*-*ε4* carriers exhibit earlier onset of β-amyloidosis relative to other alleles (21); animal studies have suggested that earlier onset of deposition can be attributed to impaired ability of apoE4 to clear soluble forms of Aβ peptide early in life (22). Other deleterious phenotypes have been attributed to *APOE4* in mice, including altered synaptic integrity and increased neuroinflammation, some of which may be amyloid-independent (23). In light of work tying the systemic environment to CNS health (6), the relative contribution of peripheral and CNS apoE pools in mediating risk is poorly understood. Recent studies highlight the role of hepatic apoE expression in regulating synaptic integrity in mice with humanized *APOE4* livers (24, 25), while ablation of hepatic apoE on its own failed to alter cerebral amyloid deposition (26). In cases of symptomatic disease or advancing age, specific proteins may be altered in cerebrospinal fluid (CSF) (27-29) and plasma of *APOE4-*carriers (29-33). It is thus likely that peripheral apoE regulates the systemic environment in an allele-dependent manner, though the characterization of how *APOE4* affects the plasma proteome in cognitively normal subjects is lacking.

While promising advances have been made that compare differential protein abundance in tissue (34) or CSF (27, 28) that provide a readout of brain health, sensitive methods are needed to probe subtle changes conferred by AD risk variants and in a manner that minimizes discomfort to subjects. For these reasons, a growing body of work has explored the use of minimally invasive peripheral markers, namely blood-based markers associated with AD signatures like amyloid-β deposition (35-41). Blood-based measurements allow for relatively convenient sampling, often before symptomatic disease onset. Conventional methods to measure plasma proteins suffer from high technical variability, or there is a lack of suitable immunoassays. To overcome these issues, we used the SOMAscan assay, which utilizes slow off-rate modified aptamers (SOMAmers) to sensitively provide relative quantification of proteins in human plasma (30, 42-48) with exceptionally low technical variation (1-4% coefficient of variation), allowing for reduced cohort sizes (49). In our plasma proteomic analysis, the SOMAscan assay measured 1,305 proteins in 53 elderly subjects homozygous for *APOE3* or *APOE4* and free of cognitive impairment. These measurements facilitated identification of blood-borne proteins associated with possession of *APOE-ε4* alleles, as well as those associated with brain amyloid positivity, regardless of *APOE* status. We performed a series of ontology-based analyses to highlight molecular markers and pathways in the blood that differed according to *APOE* or amyloid status. Specifically, we found that expression of *APOE4* leads to plasma protein perturbations associated with altered atherosclerosis, lipid transporter, ECM, and synaptogenesis signaling pathways. In contrast, presence of brain amyloid was associated with plasma protein disruptions linked to immunogenic cell death, blood-brain barrier function, and pyroptosis signaling pathways. Our findings, coupled with accumulating evidence that blood-borne proteins modulate CNS health (6), support the concept that the disruption present within the systemic environment in these contexts may represent putative targets for clinical intervention.

## Methods

### Plasma Collection

This study was performed in accordance with all appropriate human protections procedures; samples were provided as de-identified samples from the Charles F. and Joanne Knight Alzheimer Disease Research Center (ADRC) at Washington University School of Medicine, and our study received an IRB human research exempt determination from Icahn School of Medicine at Mount Sinai to carry out the research. Following recruitment, *APOE* allelic status was determined by genotyping rs7412 and rs429358 using Taqman genotyping technology (50). Clinical Dementia Rating (CDR) scores were assigned following neurological examination according to established protocols (51). Whole blood samples were drawn from fasted subjects into 10 mL syringes pre-coated with 0.5 M EDTA and transferred to 15 mL polypropylene tubes containing 120 μL of 0.5 M EDTA. Samples were kept on ice until centrifugation (<2 hours) to separate plasma from the cellular fraction. Plasma was then pooled and aliquoted into new polypropylene tubes prior to storage at -80°C. Brain amyloid status was assigned using a threshold based on previous work for CSF Aβ42 <1,098 pg/mL (amyloid-positive) or CSF Aβ42 >1,098 pg/mL (amyloid-negative) determined by the Elecsys CSF Aβ42 assay (37, 52). Male and female subjects with CDR=0 who were homozygous for either *APOE3* or *APOE4* were selected for plasma proteomic measurements.

### SOMAscan Assay

Proteomic profiling of the 1,305 SOMAmers were performed using the 1.3k SOMAscan Assay (42) at the Genome Technology Access Center (WashU) as fee-for-service. The SOMAscan plate is designed to include buffer wells and SomaLogic quality control and calibrator samples, with all samples run in duplicate on the plate. The 12 hybridization elution controls were removed prior to final analyses. Data were reported as SOMAmer reagent abundance in relative fluorescence units (RFU) with reagent abundance indicating protein concentration within the sample. Normalization of data was conducted as follows: raw data were processed for hybridization normalization to adjust for individual sample variance. Median signal normalization was then performed to adjust sample-to-sample differences in RFU brightness. Calibration normalization was then applied to all samples in the plate to remove variance due to assay run according to manufacturer recommendations.

### Covariate Analysis

The variancePartition (53) package in R (version 4.1.2) was used to calculate the proportion of variance in protein abundance explained by *APOE* genotype, amyloid status, age, sex, and SOMAscan subarray. The SOMAscan subarray was included to account for technical variability in protein abundance in the assay. The subarray is a concatenated variable that takes into account sample location (row) and plate number. The variancePartition package uses linear mixed model-based assessment to quantify the source of variation attributed to each variable and guide covariate selection for final analyses. The “canCorPairs” function was used to further quantify and interpret covariate drivers of variance.

### Protein Differential Expression Analysis (DEA)

The Differential expression for repeated measures (Dream) (54) R package was used to test the association of proteins with either *APOE* genotype or brain amyloid status. When testing differential abundance of proteins according to *APOE* genotype, only amyloid-positive *APOE3* and *APOE4* subjects were included, as amyloid-negative *APOE4* subjects were not available for cohort selection. When evaluating the effects of brain amyloid status (i.e., amyloid-positive versus amyloid-negative) on protein DEA, *APOE4* subjects were excluded for the same reason. Age, sex, and the SOMAscan subarray were covariates for such analyses. SOMAscan’s 12 elution proteins included in the assay were removed and the remaining 1,305 proteins were analyzed. Proteins were ranked according to differential abundance using log_2_ fold changes. For pathway discovery, nominal *P* values from differential expression were used as threshold for input lists, and *P* values calculated for subsequent pathways were adjusted for multiple comparisons. Top proteins were represented using boxplots created using the R package “beeswarm” from the residualized protein outputs following Dream. Briefly, covariates for age, sex, and SOMAscan subarray were regressed out while the effects of *APOE* genotype or amyloid status were re-introduced.

### DEA visualization

Volcano plots following DEA were constructed using the R package ggplot2. Customization of these plots used the packages ggrepel, dplyr, and ggeasy. The top five proteins according to - log_10_(p-value) were labeled for both the upregulated and downregulated proteins. Z-scored values were used for unsupervised hierarchical protein cluster analysis in Cluster 3.0 before heat map visualization using Java TreeView 1.0.13 based on *P*-value score ranking of the top 50 percent of significant proteins following DEA.

### Pathway analyses

Gene ontology analyses were performed using the clusterProfiler (55) package in R. Proteins that were differentially expressed at *P* < 0.05 were included in the analysis. This protein list was then sorted and analyzed separately, with proteins classified as either upregulated or downregulated using the direction of the log_2_ fold change calculated from Dream. All 1,305 proteins measured by the SOMAscan assay were used as the “universe” to evaluate the protein lists. Visualization of the functional enrichment results were carried out by dotplots using “enrichGO” and set to show the top 10 terms according to *P*-value for the GO categories (biological processes, molecular functions, and cellular components). GO terms of interest from the respective dotplots were visualized further using the “treeplot” function and proteins from these terms were extracted and evaluated for gene-network connectivity using “cnetplot”. Pathway enrichment analysis was carried out using IPA (Qiagen). Proteins differentially expressed at *P* < 0.05 were divided according to direction of change, with upregulated proteins submitted separately from downregulated proteins. IPA knowledge base was used as the reference background with a cutoff of *P* < 0.05. The top 10 significant pathways (*P* < 0.05) adjusted for multiple comparisons were selected and visualized using GraphPad Prism v9. Protein identification analysis was performed after submission to IPA and after extracting the protein classifications from the molecules tab. Plots were created using the R package Circos (56). Additional pathway enrichment analysis was carried out using the Database for Annotation, Visualization, and Integrated Discovery (DAVID; v2022q3). Briefly, differentially expressed proteins were included as the “protein list”, and all proteins measured by the SOMAscan assay were used as “background” list. Upregulated and downregulated proteins were separately analyzed, and the enrichment analysis was performed for the Reactome Pathway database through DAVID. The top 10 pathways (*P* < 0.05) were visualized and reported using the fold enrichment values calculated by DAVID.

### Weighted Gene Correlation Network Analysis (WGCNA)

Scale-free co-expression networks were constructed from proteins using the R package WGCNA (57) to identify protein modules with coordinated expression patterns according to *APOE* genotype or amyloid status. An adjacency matrix was calculated using soft powers at which the scale-free topology fit index reached 0.90 with a mean connectivity near 0. The adjacency matrix was then transformed into a topological overlap matrix (TOM). We performed this analysis using the default parameters for signed networks with exception for the following settings: soft threshold power = 14, minimum module size = 10, cutting height = 0.99, merge cut height = 0.25, and deepSplit = TRUE. All 1,305 proteins from the SOMAscan assay were assigned to modules represented by distinct color identifiers. Proteins that did not meet the criteria for module assignment were collectively placed in the “grey” module. The resulting modules of co-expressed proteins were then used to calculate module eigenproteins. Modules with a significance level of *P* < 0.05 were included in downstream analysis by gene ontology. Proteins from significant modules were extracted to identify protein-protein interactions using nodes and edges generated by the “exportNetworkToCytoscape” function and then visualized using the ggraph R package.

## Results

### Variation in *APOE* genotype alters the plasma proteome

To begin to characterize changes in plasma protein levels due to variation of *APOE3* vs. *APOE4* alleles, we measured 1,305 proteins in blood plasma from 36 *APOE3/3* and 17 *APOE4/4* elderly subjects (mean age = 68 years) (**Figure 1A-B**), all of whom were cognitively normal, according to CDR scoring (**Figure 1B**). We identified significant upregulation and downregulation in many plasma proteins associated with expressing *APOE4* relative to *APOE3* (**Figure 2A**). Unsupervised hierarchical clustering revealed distinct separation according to *APOE* status in terms of plasma proteins (**Figure 2B**). To identify the contribution of possible sources of variation within the dataset, we performed variance partition analysis (53). This analysis revealed that much of the variation is attributed to unknown biological or technical variables (**Figure 2C**). To further decouple sources of variation, we also performed a canonical correlation analysis to identify the degree with which variables co-vary. Findings from this analysis suggest correlation between *APOE* genotype and other variables, including brain amyloid status (**Figure 2D**). Thus, while *APOE4* alters the plasma proteome relative to expression of *APOE3* in elderly subjects, controlling for additional sources of biological and technical variation may improve accuracy in assessing individual protein-level changes conferred by *APOE4*.

**Figure 1.**
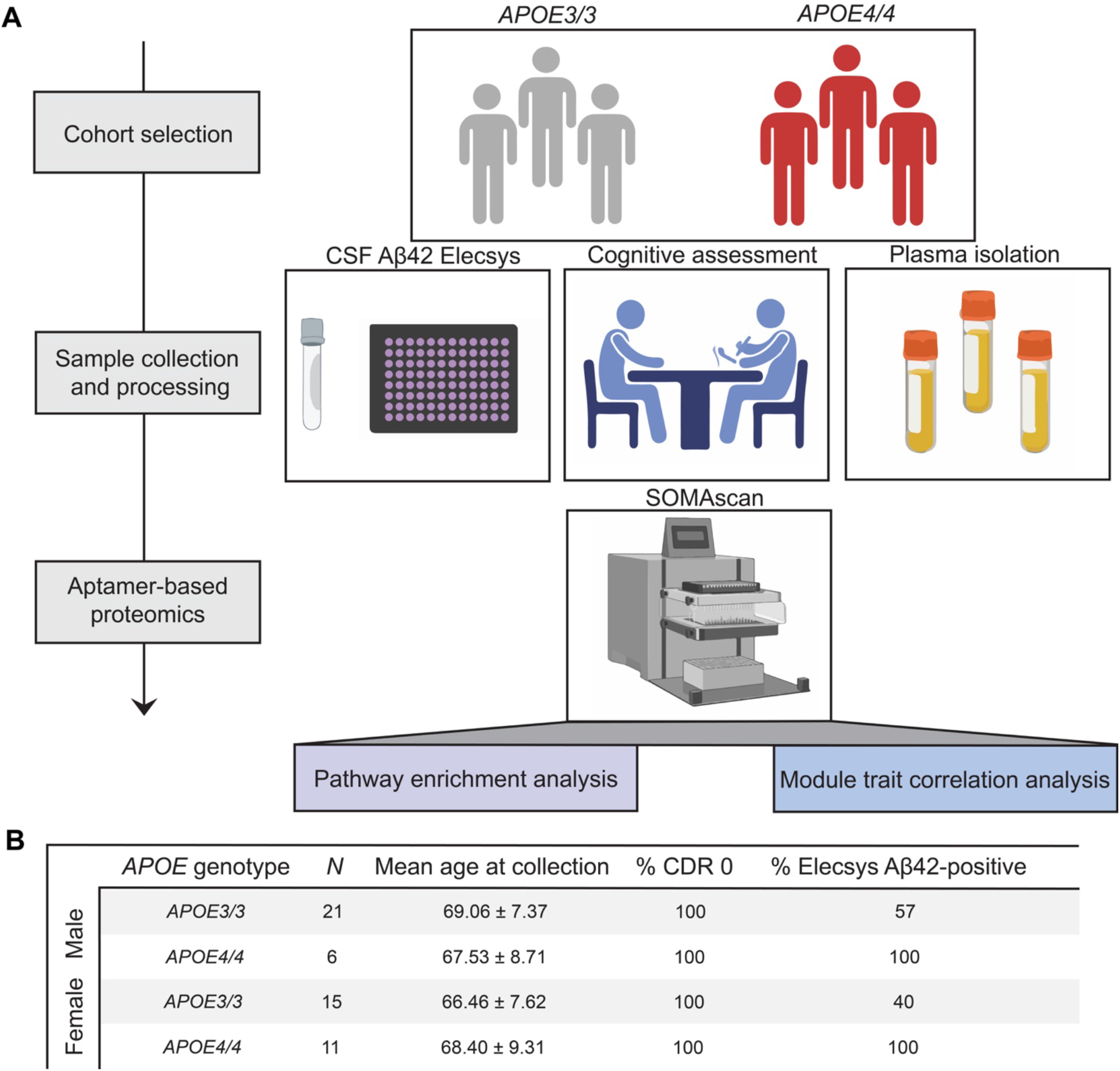
Overview of the study design and cohort demographics. **A**. Schematic workflow describing selection and measurement of plasma samples from 53 *APOE3* and *APOE4* subjects from the Knight Alzheimer’s Disease Research Center (WashU). CSF samples were collected from subjects by the Knight ADRC and run on Aβ42 Elecsys assays. Subjects underwent neurological examination and were classified according to clinical dementia rating (CDR). Fresh blood samples and isolated plasma from *APOE3* and *APOE4* subjects were collected following established protocols. Relative abundance of 1,305 proteins was measured using SOMAscan aptamer-based profiling, followed by examination of *APOE* and amyloid-associated effects on pathway enrichment and module-trait relationships. **B**. Male and female subjects with homozygous expression for *APOE3* or *APOE4* were evaluated. All participants had CDR of 0. *APOE3* subjects were either amyloid-positive or amyloid-negative, whereas all *APOE4* subjects were amyloid-positive, due to subject availability.

**Figure 2.**
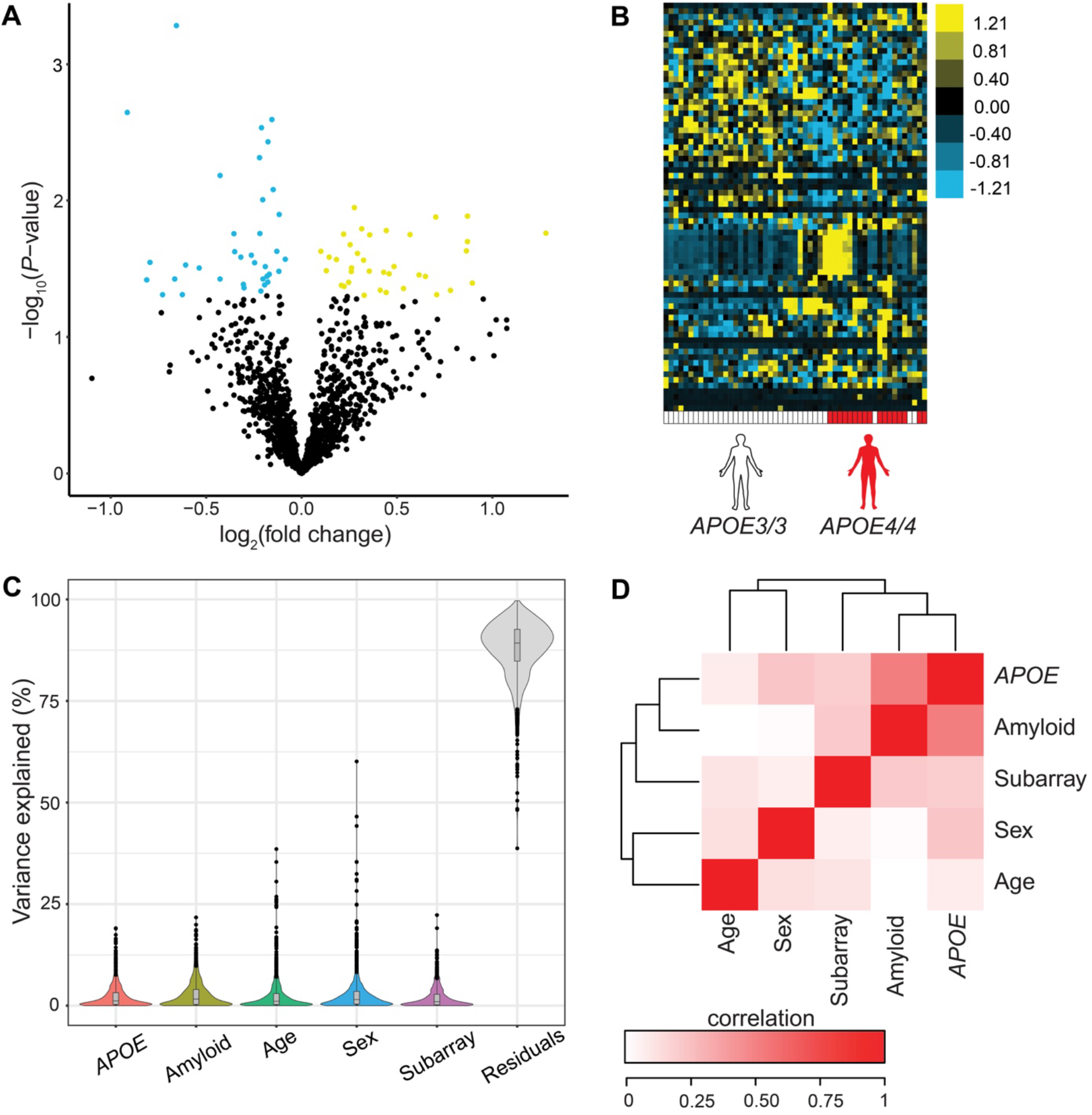
Variation in *APOE* alters the plasma proteome in elderly adults. **A** Volcano plot showing fold-change of proteins (log_2_ scale) between *APOE3* and *APOE4* subjects and corresponding nominal *P*-values (-log_10_ scale) with upregulated proteins highlighted in yellow and downregulated proteins highlighted in blue. **B**. Heat map showing unsupervised hierarchical clustering of the differentially abundant proteins distinguished by *APOE* status. **C**. Variance partition plot showing variance attributed to certain covariates. **D**. Covariate correlation analysis of biological and technical sources of variability. Degree to which these variables co-vary is highlighted from white (no level of co-variance) to red (high degree of co-variance).

### *APOE* genotype distinguishes plasma proteomic changes after controlling for covariates

To evaluate downstream differences in plasma protein abundance associated with *APOE4* expression, we performed a linear mixed effects regression analysis using the Dream package (54) and controlled for both biological and technical variability. We identified *APOE4*-associated changes in 80 plasma proteins, with several proteins significantly upregulated or downregulated, even after adjusting for these variables (**Figure 3A**). Unsupervised hierarchical clustering revealed distinct separation of plasma protein abundance between *APOE3* and *APOE4* subjects (**Figure 3B**). We found a significant increase in abundance for 39 proteins and a decrease in abundance for 41 proteins between *APOE4* and *APOE3* subjects (**Figure 3C**), with sizeable proportions belonging to diverse protein categories, as visualized using protein-type identification analysis. This confirms our use of an unbiased platform (42) and the tendency of the *APOE-ε4* allele to associate with broad changes in the plasma proteome. The top ten differentially expressed proteins included NME2, UNC5D, EFNB2, CFP, and PPBP, as well as SLAMF7, CRP, AURKA, FUT5, PAK6, which were up- and downregulated, respectively, in *APOE4* relative to *APOE3* subjects (**Figure 3D**). Interestingly, NME2, which is significantly increased in *APOE4* subjects, is an important regulator of the calcium-activated K+ channel K_Ca_3.1. These channels play a role in mediating CD4+ T cell activation (58), a process recent work has suggested may be dysregulated in AD (59, 60). We also found that SLAMF7 (**Figure 3E**) was the top downregulated protein associated with *APOE4*. SLAMF7 has been implicated as a key regulator of adaptive immunity (61), which may regulate AD pathogenesis (62). Together, these data indicate that *APOE* allelic variation can significantly alter plasma proteins previously implicated in AD prior to symptom onset and in a manner unrelated to amyloid status.

**Figure 3.**
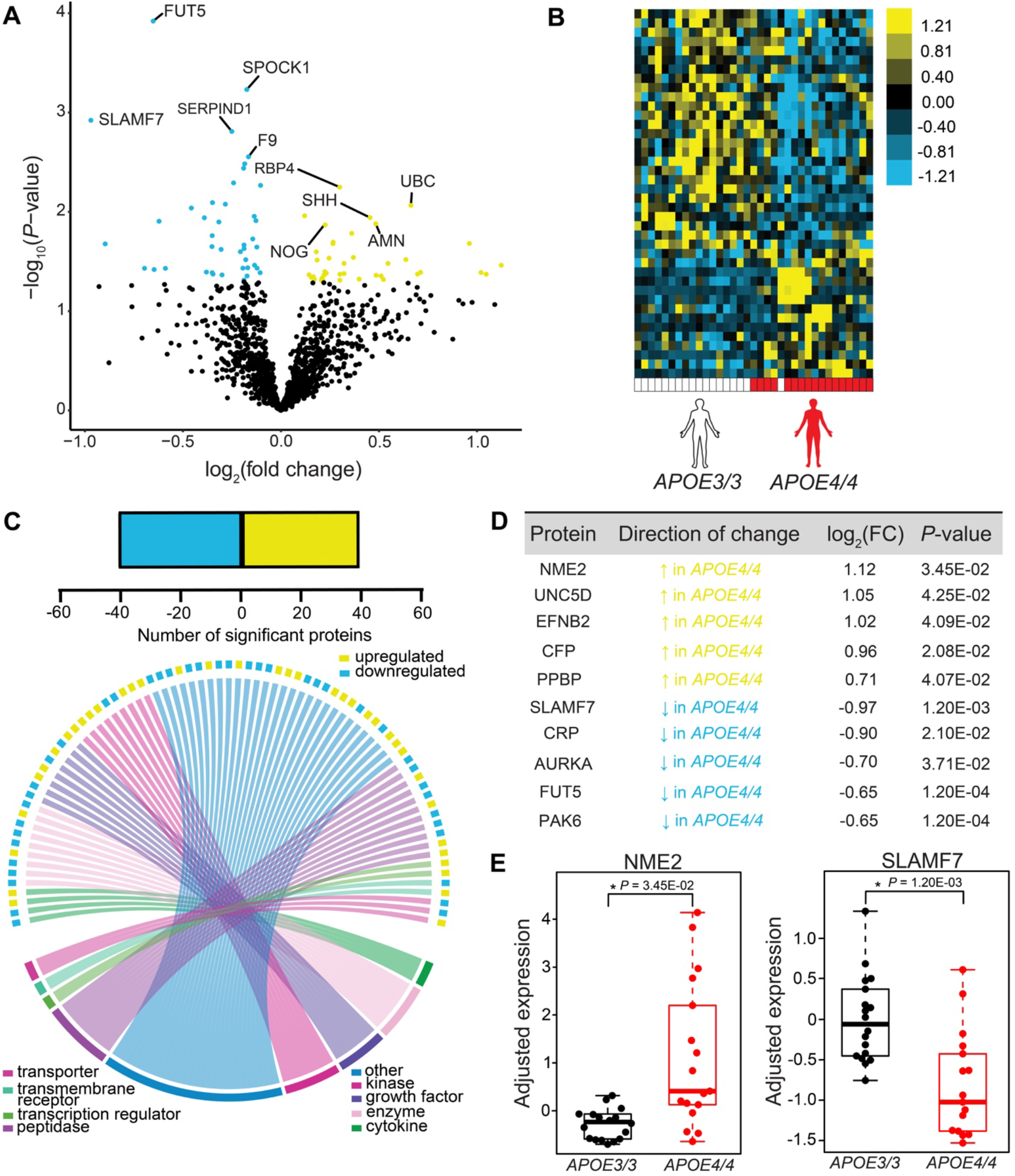
Plasma proteome features from cognitively normal, amyloid-positive *APOE3* and *APOE4* subjects. **A**. Volcano plot showing the fold change of proteins (log_2_ scale) between *APOE3* and *APOE4* subjects and corresponding nominal *P*-values (-log_10_ scale) with upregulated proteins highlighted in yellow and downregulated proteins highlighted in blue. **B**. Heat map showing unsupervised hierarchical clustering of the differentially expressed proteins distinguished by *APOE* status following covariate adjustment using Dream package. **C**. (Upper) Number of significant up- and downregulated proteins. (Lower) Circos plot mapping of protein family types measured in differentially expressed proteins between *APOE3* and *APOE4* subjects. Individual color-coded labels indicate protein category types. **D**. Top 10 differentially expressed proteins ranked by log_2_ fold-change between *APOE3* and *APOE4* subjects following Dream package. **E**. Sample top significant proteins plotted using residualized Dream expression values. *P*-values were calculated with Dream.

### *APOE4* alters pathways in plasma proteome associated with inflammation and CNS function

To evaluate whether plasma proteomic pathways are dysregulated according to *APOE4* status, we then submitted these plasma proteins with differential abundance to Ingenuity Pathway Analysis (IPA). This analysis revealed several significant pathways associated with immune function, including STAT3 and IL-15 production, as well as predicted changes in atherosclerosis signaling (**Figure 4A**). Surprisingly, brain-specific pathways were also identified from *APOE4*-driven plasma protein changes, including changes in “synaptogenesis” and “axonal guidance” (**Figure 4A**). To investigate metabolic processes associated with the *APOE-ε4* allele, we also analyzed pathways from the Reactome database, which is preferentially modeled using biological reaction data (63). We submitted upregulated proteins associated with *APOE4* to DAVID and analyzed significantly altered pathways (*P* < 0.05) from the Reactome database, which revealed changes in “plasma lipoprotein clearance” and “diseases of metabolism” (**Figure 4B**), as expected (64). Interestingly, we again saw brain-specific pathways like “axon guidance” and “nervous system development”, reinforcing CNS-related pathways identified earlier by IPA. Together, these analyses suggest that expression of *APOE4* can alter pathways important within the periphery and central nervous system. We next sought to explore putative functional changes associated with *APOE4* in plasma by evaluating gene ontology terms enriched by *APOE4*-associated plasma proteins (**Figure 4C**). Similar to our pathway expression analysis, we again found expected functions known to be associated with the *APOE-ε4* allele, including “lipid transporter activity” and general “transporter activity”. We also found molecular function terms related to the extracellular matrix (ECM), including “ECM” and “laminin binding”. Given that previous work has implicated the ECM as an important mediator in blood-CNS communication (7), we sought to further probe the underlying plasma proteins associated with these functions and their relatedness. To do this, we constructed a hierarchical treeplot, which indicated a close relationship between ECM and laminin binding downstream of known *APOE4*-associated lipid and transporter activities (**Figure 4D**). After extracting proteins related to these terms, we created a gene-concept network plot (**Figure 4E**) and found plasma proteins were specific to either the ECM (LGALS3 and SHH) or related to transporter activity terms (RBP4 and PPBP). Together, our results suggest that expression of *APOE4* may induce changes to the ECM that may be related to, but separate from, known apoE protein-specific functions, such as lipid transporter activity.

**Figure 4.**
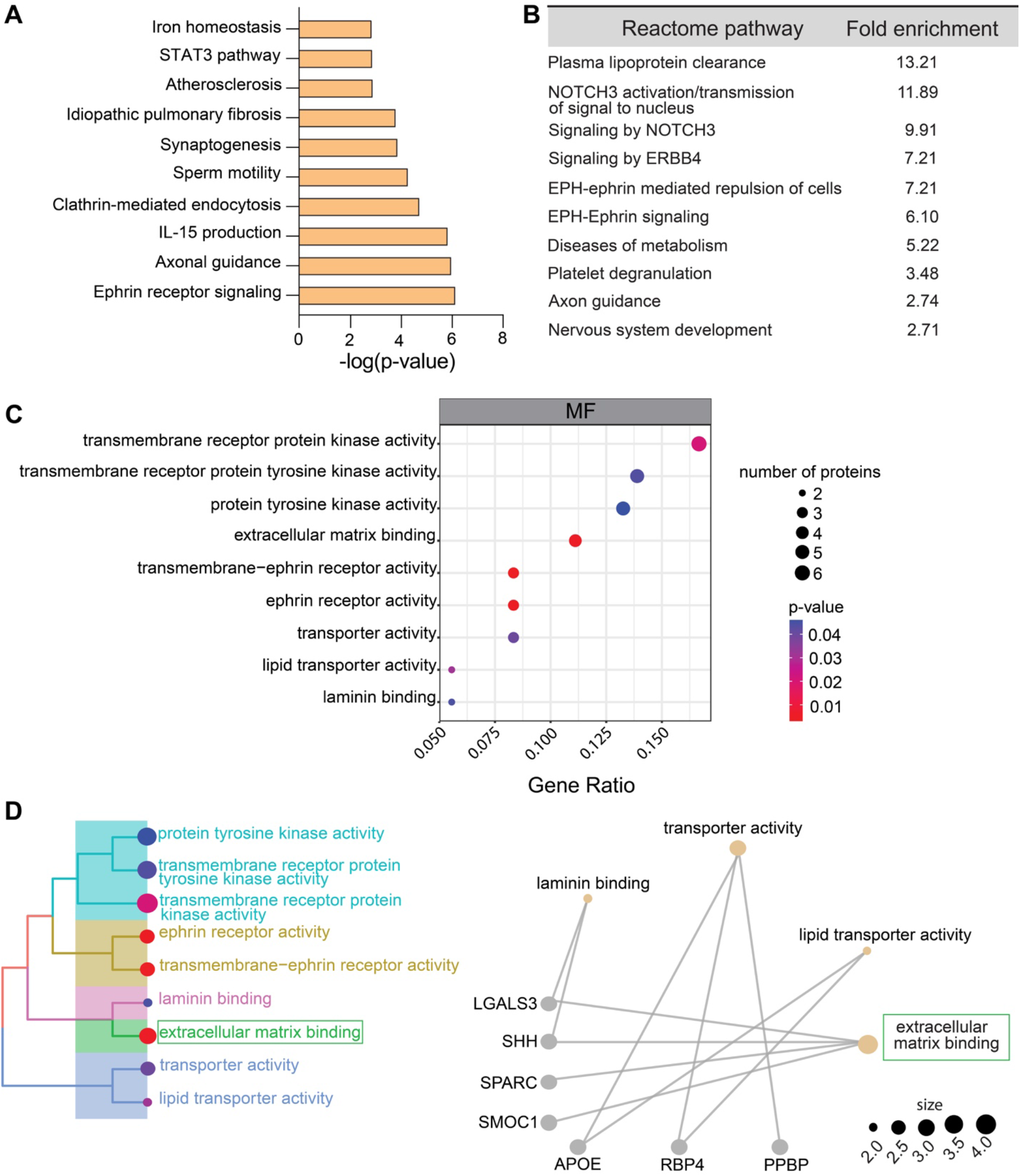
*APOE4* expression alters plasma protein pathways and networks. **A**. Pathway analysis from IPA for plasma proteins upregulated by *APOE4*. Significance is represented by -log(p-value) for the top 10 pathways. **B**. Pathway enrichment analysis using the Reactome Database from DAVID. The top 10 most significant pathways (*P* < 0.05) were shown with corresponding fold-enrichment values, calculated as proportion of input proteins and total number of proteins associated with that pathway. **C**. Enrichment analyses were performed using *APOE4*-associated proteins for Gene Ontology molecular functions (MF). MF terms highlighted were chosen according to *P*-value with color indicating significance and size indicating number of proteins. **D**. MF terms were further analyzed by a hierarchical treeplot, where distance on the plot indicates similarity, color indicates significance, and size indicates number of proteins. Visualization of these MF terms by gene network plots was used for further investigation.

### Co-regulatory protein expression changes are induced in *APOE4* subjects

To examine how plasma proteins are co-expressed and co-regulated in the *APOE4* plasma proteome relative to *APOE3*, we used a weighted gene co-expression network analysis (WGCNA) (**Supplementary Figure 1A-D**), which revealed a significant module (cyan) that was positively correlated with *APOE4* status (*P* = 3.22 × 10^−2^) (**Supplementary Figure 2A-B**). After extracting genes from the cyan module, we identified mapping of several hub proteins (**Supplementary Figure 2C**). These hub proteins were found to be related to ECM activity (SARC, THBS1, PDGFB) and immune/inflammatory responses (PFA and PPBP). To identify the ability of these hub proteins to influence functional analyses of the cyan module broadly, we performed a Gene Ontology enrichment analysis on cyan module proteins (**Supplementary Figure 2D**). We identified biological processes related to immune cell movement and migration (leukocyte migration and leukocyte chemotaxis), while cellular components and molecular functions were related to the ECM (collagen-containing ECM, basement membrane, and collagen binding). This suggests that altered immune function and ECM regulation may be critical mediators of broad *APOE4*-mediated regulatory changes within the plasma proteome.

### Impact of brain amyloid on plasma proteins in *APOE3* subjects

We next sought to examine whether brain amyloid-positivity itself was associated with altered plasma protein abundance independent of *APOE4* status, which might imply disease-specific peripheral dysregulation. We examined relative protein abundance by SomaScan from 36 *APOE3* subjects, 18 of whom were categorized as amyloid-positive, while the other 18 were amyloid-negative (**Figure 1B**). *APOE4* subjects were not included given that sufficient numbers of amyloid-negative *APOE4* subjects were not available for comparison. We applied the Dream model to adjust covariates and then created a volcano plot to visualize significantly upregulated or downregulated proteins driven by presence of brain amyloid (**Figure 5A**). Unsupervised hierarchical clustering revealed a distinct plasma protein profile between amyloid-positive and amyloid-negative subjects (**Figure 5B**). Notably, we found that abundance of 91 proteins was significantly increased, while abundance of 61 proteins was significantly decreased in amyloid-positive relative to amyloid-negative subjects (**Figure 5C**), all of which were well represented across protein classes. The top 10 differentially expressed proteins distinguished by presence of brain amyloid were SAA1, CSNK2A1, CD33, PAK6, HIST2H2BE, as well as NME2, UNC5D, EFNB2, NAAA, MICA, which were up- and downregulated, respectively. Interestingly, levels of NME2 in amyloid-negative *APOE3* subjects are elevated (**Figure 5D-E**), likely driven by the low levels of this protein in amyloid-positive *APOE3* subjects also observed in our previous comparison (**Figure 3D**). Notably, CD33, a top AD risk gene linked to microglial function that was identified through genome-wide association studies (65-67), was upregulated in plasma in amyloid-positive subjects (**Figure 5D**). We also found that SAA1, a protein known to be highly abundant in AD cases vs. controls (68, 69), was the top elevated plasma protein in amyloid-positive vs. amyloid-negative subjects (**Figure 5E**).

**Figure 5.**
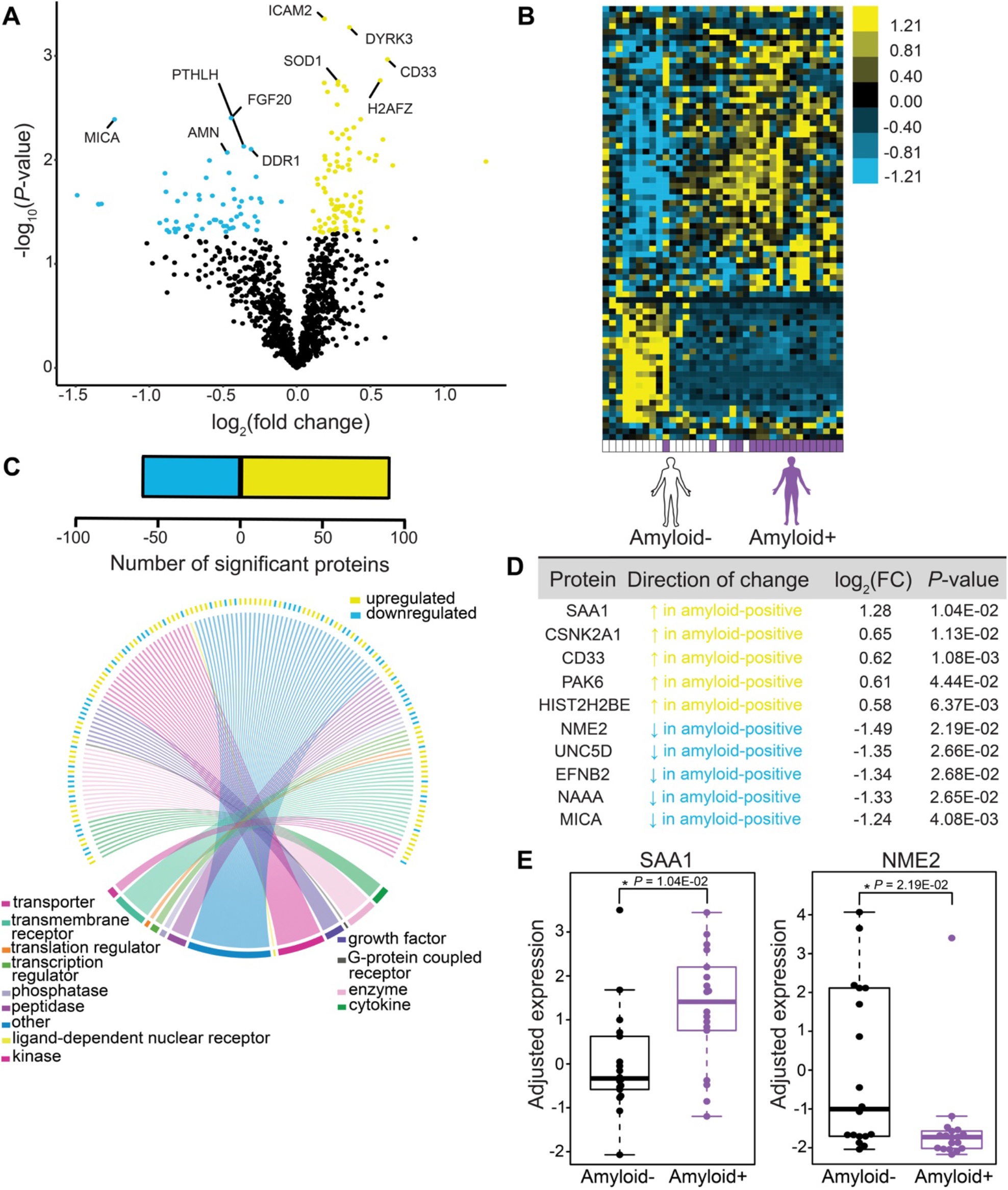
Presence of brain amyloid alters plasma proteome features in *APOE3* subjects. **A**. Volcano plot showing the fold change of proteins (log_2_ scale) between amyloid-positive and amyloid-negative subjects and corresponding nominal *P*-values (-log_10_ scale) with upregulated proteins highlighted in yellow and downregulated proteins highlighted in blue. **B**. Heat map showing unsupervised hierarchical clustering of the differentially expressed proteins distinguished by brain amyloid status following covariate adjustment using Dream package. **C**. (Upper) Number of significant up- and downregulated proteins. (Lower) Circos plot mapping of protein family types measured in differentially expressed proteins between *APOE3* subjects who are amyloid-positive or amyloid-negative. Individual color-coded labels indicate protein category types. **D**. Top 10 differentially expressed proteins ranked by log_2_ fold-change between *APOE3* amyloid-positive and amyloid-negative subjects following Dream package. **E**. Sample top significant proteins plotted using residualized Dream expression values. *P*-values were calculated with Dream.

### Brain amyloid alters pathway and network function in plasma of *APOE3* subjects

To examine the pathways in the plasma proteome disrupted by the presence of brain amyloid, we submitted differentially expressed plasma proteins associated with amyloid positivity to IPA (**Figure 6A**). Pathways related to altered immune function and inflammation observed in AD subjects (70) were identified, including “immunogenic cell death”, “leukocyte extravasation”, and “IL-15 production”. Further probing of pathways significantly altered by amyloid-positivity was conducted using the Reactome database (**Figure 6B**), which revealed pathways related to “DNA methylation” and “chromatin organization”. Additionally, a “defective pyroptosis” pathway was identified, possibly indicating processes linked to increased cell death (71). We probed this further using Gene Ontology enrichment analysis on the plasma proteins associated with amyloid-positivity and found consensus with altered biological processes related to chromatin organization and assembly (**Figure 6C**). Notably, we found several CNS-associated biological processes associated with brain amyloid that were linked to plasma proteomic changes, including “axon and neuronal ensheathment”, “myelination”, and “maintenance of blood-brain barrier”. Given previous links between AD and blood-brain barrier (BBB) disruption (72, 73), we explored the plasma proteins underlying this association and its related terms by constructing a hierarchical treeplot. “Maintenance of the blood-brain barrier” was found closely associated with functions related to protein localization to “cell junctions” and “cell-cell junction assembly” (**Figure 6D**). As expected, overlapping plasma proteins identified by gene-network construction were related to cellular adhesion, including JAM2, JAM3, CDH5, PECAM1 (**Figure 6E**). Our results point to plasma protein changes associated with brain amyloid that are linked to BBB maintenance in cognitively normal subjects.

**Figure 6.**
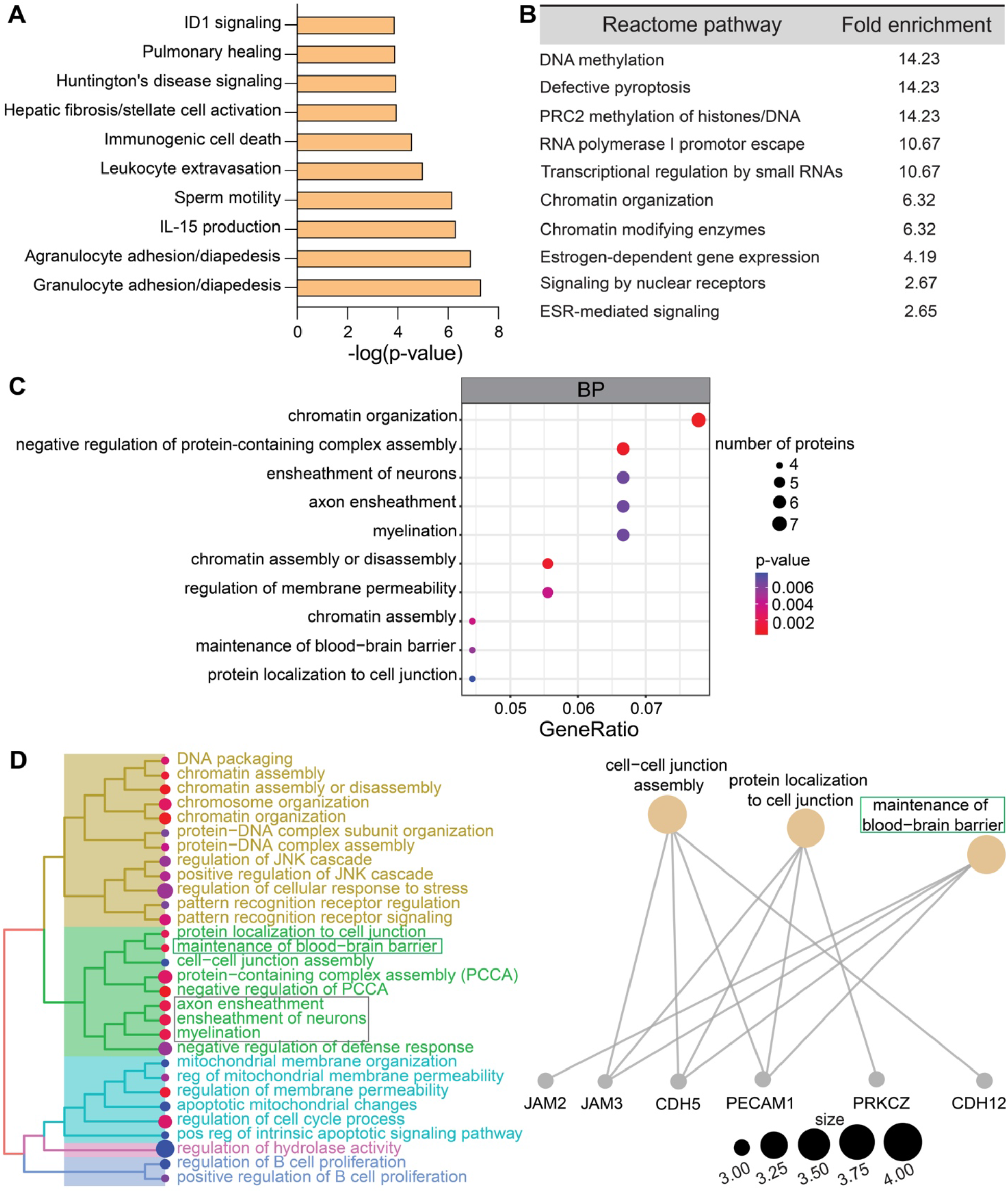
Brain amyloid-positivity alters pathway and network annotations. **A**. Pathway analysis from IPA for plasma protein upregulated by brain amyloid positivity. Significance is represented by - log(p-value) for the top 10 pathways. **B**. Pathway enrichment analysis using the Reactome Database from DAVID. Top 10 most significant pathways (*P* < 0.05) were shown with corresponding fold-enrichment values, calculated as proportion of input proteins and total number of proteins associated with that pathway. **C**. Enrichment analyses were performed using amyloid-associated proteins for Gene Ontology biological processes (BP). BP terms highlighted were chosen according to *P*-value, with color indicating significance and size indicating the number of proteins. **D**. BP terms were further analyzed by a hierarchical treeplot where distance on the plot indicates similarity, color indicates significance, and size indicates the number of proteins. Visualization of these BP terms by gene network plots was used for further investigation.

### Co-regulatory protein expression changes are induced in amyloid-positive subjects

To further understand how plasma proteins altered in the context of brain amyloid are co-regulated, we employed WGCNA and found significant positive upregulation of the brown module (*P* = 6.87 × 10^−3^) (**Supplementary Figure 4A-B**). Hub proteins (**Supplementary Figure 4C**) mapped to the brown module were predominantly related to immunity/inflammation (CSF1, PECAM1, FCN1), cell adhesion (CDH12), and ECM regulation (MMP17, DDR2), consistent with our pathway analyses. To support the relationships of these proteins with specific functions, Gene Ontology enrichment analysis (**Supplementary Figure 4D**) identified molecular functions related to ECM function (“heparin binding” and “glycosaminoglycan binding”) and several molecular functions and cellular components related to inflammation (“chemokine activity”, “chemokine binding”, “azurophil granule lumen”, “primary lysosome”). Together, our results suggest that harboring brain amyloid is associated with distinct changes in the plasma proteome that are linked to altered immune and ECM processes and blood-borne processes linked to BBB and CNS function.

## Discussion

Given the emerging interest in characterizing the extent to which dysregulation in the periphery reflects or contributes to neurodegenerative diseases, our goal in the current study was to characterize how variation in *APOE* alters the plasma proteome in cognitively normal, elderly subjects. Given that presence of amyloid was a significant source of variation, we compared plasma proteins in amyloid-positive *APOE3* subjects to amyloid-positive subjects expressing the high risk *APOE-ε4* alleles. We found that variation of *APOE* genotype influences levels of various plasma proteins, consistent with earlier reports in *APOE4*-carriers vs. non-carriers for a subset of proteins (30). We found significant alterations associated with *APOE* status in proteins related to immune function, including in NME2 and SLAMF7, which are implicated in downstream activation of CD4+ T cells and adaptive immunity, respectively. Accumulating evidence has demonstrated a significant relationship between *APOE* and innate immune function in AD (74, 75), but the relationship between *APOE4* and adaptive immunity is less understood. Recent work has found that adaptive immunity is dysregulated in *APOE4*-carriers, with decreased abundance of C-reactive protein (CRP) relative to *APOE3* subjects, as assessed in CSF (76) and following both acute (77) and chronic (78) measurements of CRP in blood serum. Chronically low levels of CRP were also associated with accelerated AD onset (78). These studies are consistent with our findings that CRP levels are significantly decreased in *APOE4* subjects relative to *APOE3* in cognitively normal subjects (**Figure 3C**). From pathway analyses, we found that *APOE4*-associated changes in plasma were consistent with perturbations in lipid transporter activity and ECM organization **(Figure 3C-D)**, findings that were highlighted throughout Gene Ontology enrichment analyses. *APOE4*-associated alterations in lipid transporter activity are consistent with the canonical function of apoE protein throughout the body and the impact of its polymorphisms (64). Though an increase in number of subjects and protein aptamer coverage might further validate a role for *APOE4* in altering ECM function in the blood, further analysis of broad protein relationships by weighted gene co-expression analysis revealed significant co-regulation of *APOE4*-associated proteins related to the ECM. Our recent work examining age-associated human and mouse plasma proteins and their impact in blood-brain communication also implicated proteins that regulate the ECM (7). Moreover, the relationship between *APOE4* and altered ECM function has been recently suggested by several groups in the brain (79, 80). These groups have shown similar patterns related to ECM organization and regulation as we now observe, yet we now identify these changes in the systemic environment of cognitively normal, elderly subjects. Additionally, a previous study evaluating plasma protein changes in *APOE4* subjects identified unique “dementia-associated proteins” (81). While only two of the six unique proteins identified were also measured in our study, both proteins, IGFBP2 and F10, (data not shown) were not significant in our evaluation of *APOE4*-associated plasma proteins in our cohort of cognitively normal, elderly subjects, perhaps arguing that differences might be related to disease stage. Indeed, a separate study following centenarians and their offspring identified *APOE*-associated changes in 16 proteins measured in blood serum that were capable of predicting future disease susceptibility (30).

In addition to examining how *APOE* genotype contributes to the plasma proteome, we further examined contributions of AD pathology to the proteomic composition of the systemic compartment. Using an established biomarker cut-off indicative of the presence of brain amyloid (37, 52), we examined subjects who were amyloid-positive or amyloid-negative and expressing neutral *APOE-ε3* alleles. We find that brain amyloid positivity resulted in widespread changes in plasma protein abundance. One protein in particular, NME2, was significantly upregulated in amyloid-negative *APOE3* subjects, as well as in amyloid-positive *APOE4* subjects, possibly suggesting differing regulation of NME2-K_ca_3.1 function. Regulation of Ca^2+^ signaling through *Kca3*.*1* has been attributed to microglial activation (82), and its pharmacological blockade induces anti-inflammatory effects in amyloid-bearing mice (83), possibly linking changes in NME2 abundance to documented neuroinflammatory changes in *APOE4*-carriers (74). NME2 regulation also involves synthesis of nucleoside triphosphates, lipid membrane binding, and endocytic processes (84), underlining the need to further understand putative roles in AD. Plasma levels of SAA1 were found to be upregulated in amyloid-positive subjects, which was supported by pathway analysis and enrichment of gene ontology terms linked to inflammation. Pathway analysis on plasma proteins altered in amyloid-positive subjects highlighted changes in processes previously linked to AD pathogenesis, including the blood-brain barrier (85, 86), programmed cell death by pyroptosis (71), and altered epigenetic mechanisms related to DNA and chromatin manipulations (87). A study measuring plasma proteins in AD cases and controls highlighted some of these pathways (“regulation of apoptotic process” and “extracellular matrix organization”) but also distinct pathways (29), possibly due to the more advanced ages and/or pathology in their cohort. Large-scale protein co-regulatory changes identified using WGCNA indicate broad but likely interconnected changes in ontology functions related to immunity/inflammation and ECM regulation. Comparing these pathways and functions related to protein co-regulation in our dataset to previous work (29), there is some consensus, including regulation of endothelial cell migration, changes in cytokine activity, and G-protein coupled receptor binding (**Supplementary Figure 4E**). Shared pathways between the datasets may indicate broad phenotypes in the plasma proteome that reflect the long pathological process of AD across preclinical to symptomatic stages. Several studies have focused on changes in the plasma proteome in AD cases and controls (29, 47). Our study, while discovery-oriented, uniquely highlights changes in the systemic environment in cognitively normal subjects in terms of *APOE* status and brain amyloid positivity.

## Conclusions

Our results add to an ongoing framework with which to evaluate the contribution of *APOE* allelic variation or amyloid-positivity to study the complex interactions between the systemic environment and the brain. We demonstrate that *APOE4* expression or brain amyloid positivity are associated with altered abundance of diverse classes of plasma proteins. The identified pathways and functional annotations further highlight the possibility of drug-based interventions to combat AD in diverse compartments of the body.

## List of abbreviations

AD: Alzheimer’s disease
APOE: apolipoprotein E
BBB: blood-brain barrier
CDR: clinical dementia rating
CNS: central nervous system
CRP: C-reactive protein
CSF: cerebrospinal fluid
DAVID: Database for Annotation, Visualization, and Integrated Discovery
ECM: extracellular matrix
IPA: Ingenuity Pathway Analysis
SOMA: slow off-rate modified aptamers
TOM: topological matrix overlap
WGCNA: weighted gene co-expression analysis

## Ethics declarations

J.M.C. is listed as a co-inventor on patents for treating aging-associated conditions, including the use of young plasma administration (US10688130B2) or youth-associated protein TIMP2 (US10617744B2), the latter of which is licensed to Alkahest, Inc. The remaining authors have no actual or potential conflicts of interest to declare.

## Acknowledgments

We would like to extend our gratitude to all subjects involved for volunteering their time and the clinical staff for human blood-plasma collection/coordination from the Charles F. and Joanne Knight Alzheimer Disease Research Center (ADRC) at Washington University School of Medicine (St. Louis, MO). We would also like to thank Carlos Cruchaga (WashU), Rick Perrin (WashU), Anne Fagan (WashU), and Suzanne Schindler (WashU) for advice and coordination regarding ADRC cohort selection. We would also like to thank Karen Therrien (ISMMS) for advice regarding differential expression analyses. This work was funded by a 2015 New Vision Research Investigator Award (J.M.C.), the BrightFocus Foundation (A2018213S (J.M.C)), the National Institute of Mental Health (T32MH087004 (S.M.P)), the National Institute on Aging (AG061382 (J.M.C.), AG061382-02S1 (J.M.C., S.M.P)), the Healthy Aging and Senile Dementia study (P01AG003991), the WashU Knight Alzheimer’s Disease Research Center (P30AG066444), and the Adult Children Study (P01AG026276).

## Author Contributions

S.M.P. performed data analysis and wrote the manuscript. J.M.C. collected data, designed, and supervised the research, and wrote the manuscript. K.BP and T.R. assisted with data analysis and writing of the manuscript. All authors interpreted and discussed the results.

---

**Supplementary Figure 1.**
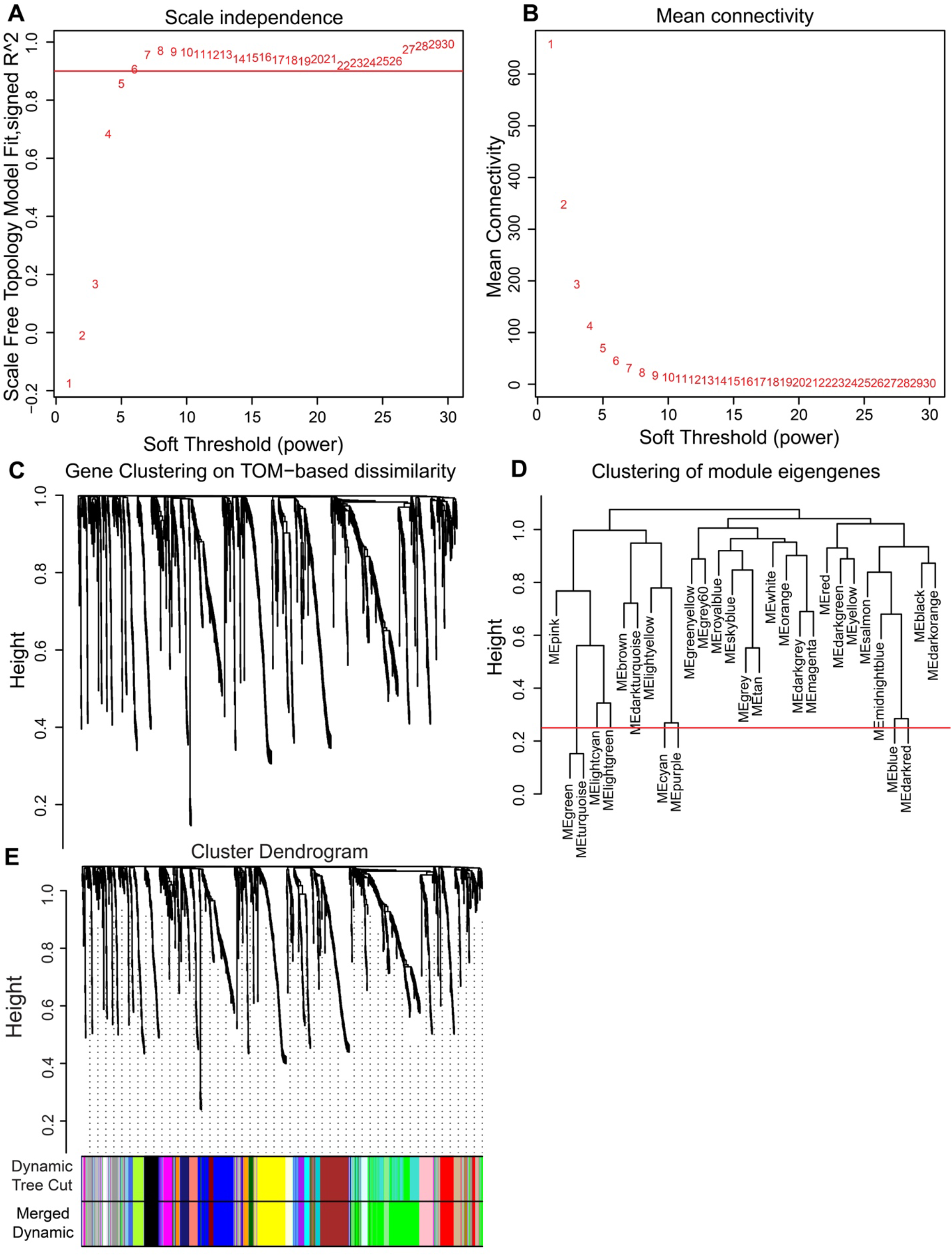
Network construction for co-regulatory protein analysis between *APOE4* and *APOE3* subjects. **A**. Network parameters were selected for soft thresholding powers. Scale-free topology is attained above the red line. **B**. Mean connectivity for further soft thresholding analysis was selected as values approach 0. **C**. Gene clustering for dissimilarity by topological overlap. **D**. Clustering of module eigengenes to identify overlapping modules was selected using a merge height of 0.25 (red line). **E**. Cluster dendrogram of topological overlap with colored bars to indicate module assignments. Dynamic tree cut shows the unmerged module assignments related to the merged dynamic for the final module colors.

**Supplementary Figure 2.**
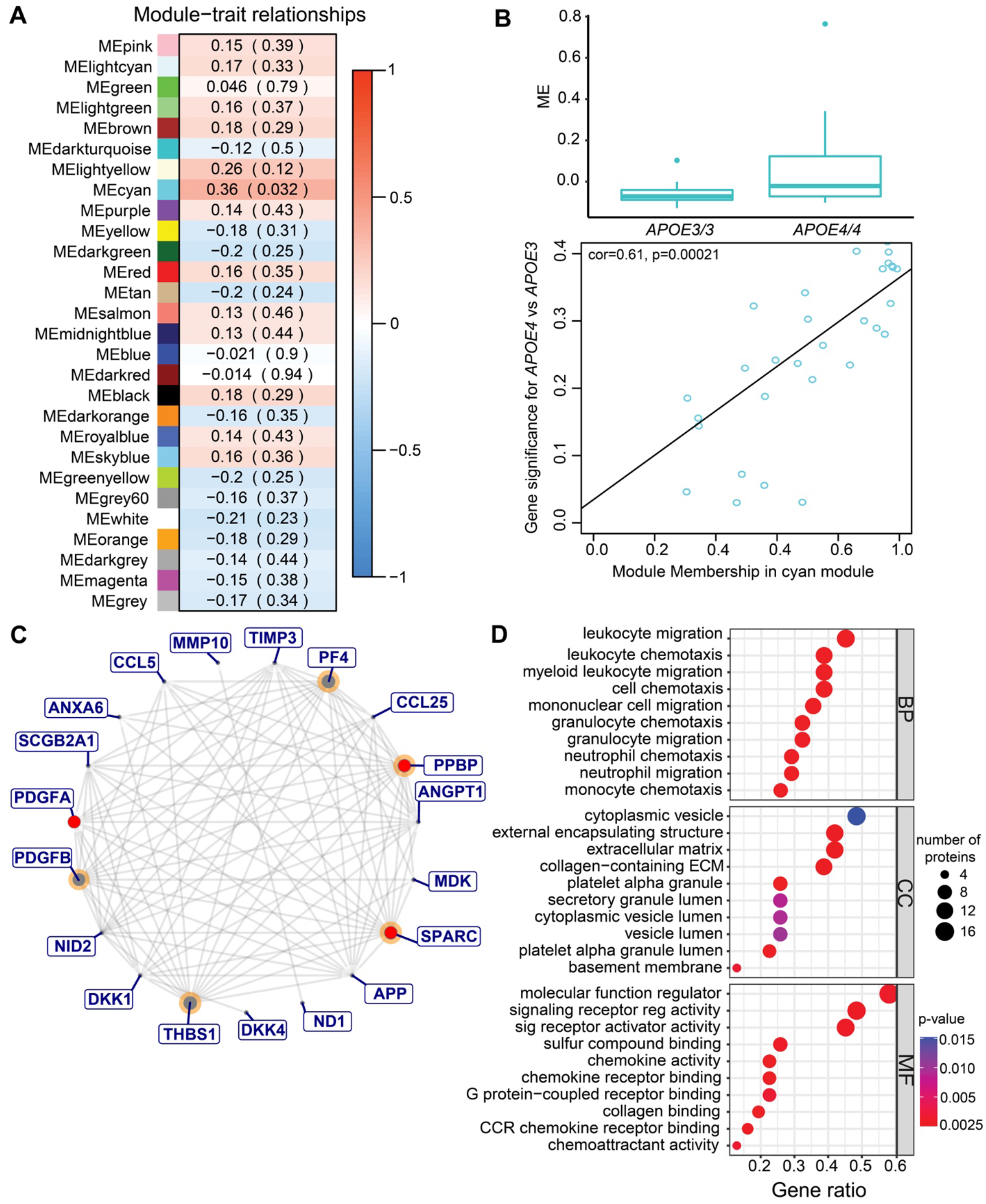
WGCNA module selection for co-regulatory changes between *APOE4* and *APOE3* subjects. **A**. Construction of the module-trait relationships plot for all module assignments (color labels), in addition to the module membership values and the respective module significance in parentheses. Selection of modules for further analysis is based on significance (*P* < 0.05). **B**. Bar graph demonstrating module eigengene differences for the cyan module between *APOE4* and *APOE3* subjects. Scatterplot indicates the correlation among individual module eigen values and comparison between *APOE* status. **C**. Extraction of proteins from the cyan module were visualized for protein-protein interactions by node and edge connectivity. Identification of hub genes (orange) was made by assessing module membership values, while significantly upregulated proteins identified previously in Dream are highlighted in red. **D**. Enrichment analyses were performed using *APOE4*-associated proteins for Gene Ontology biological processes (BP), cellular components (CC), and molecular functions (MF).

**Supplementary Figure 3.**
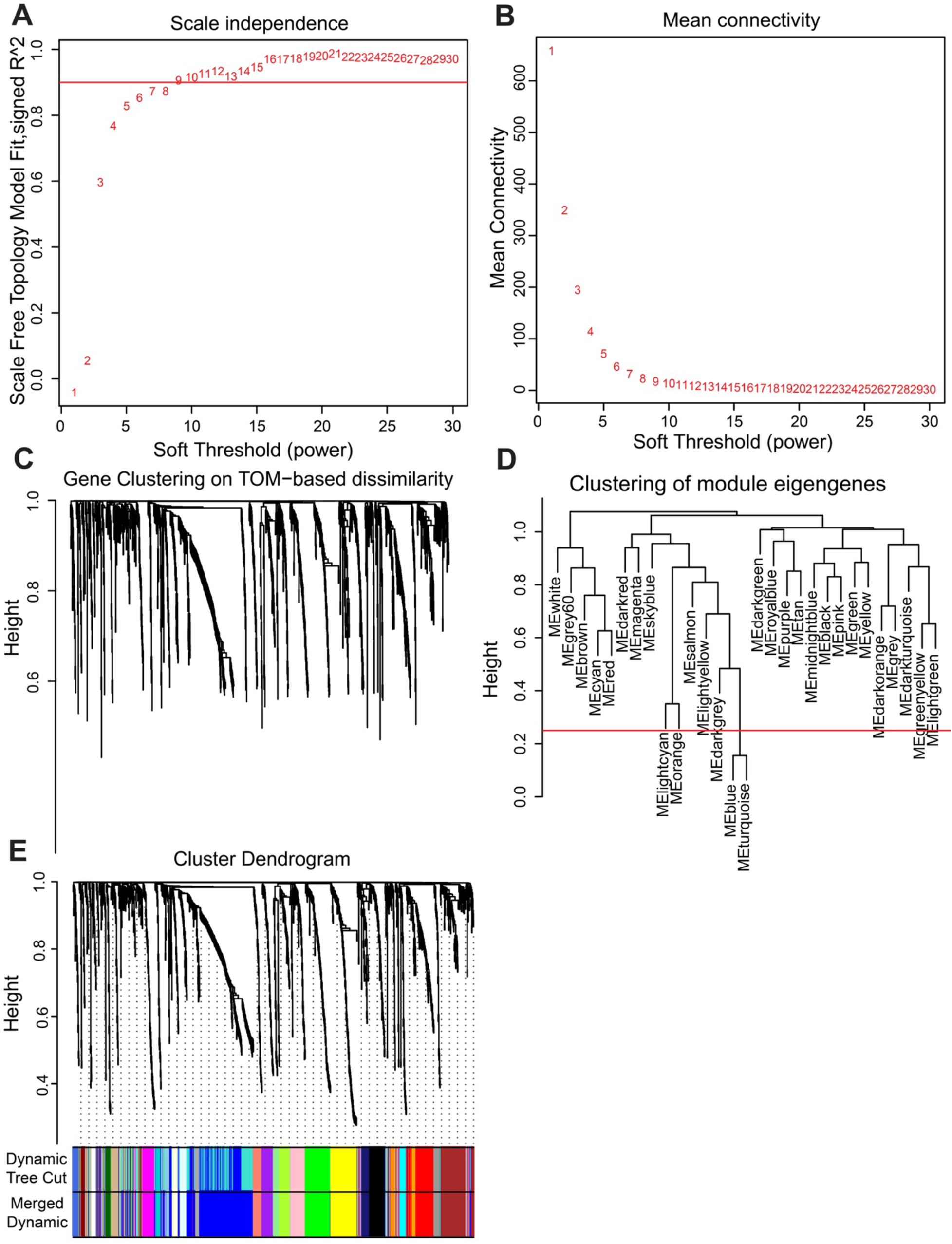
Network construction for co-regulatory protein analysis between amyloid-positive and amyloid-negative subjects. **A**. Network parameters were selected for soft thresholding powers. Scale-free topology is attained above the red line. **B**. Mean connectivity for further soft thresholding analysis was selected as values approach 0. **C**. Gene clustering for dissimilarity by topological overlap. **D**. Clustering of module eigengenes to identify overlapping modules was selected using a merge height of 0.25 (red line). **E**. Cluster dendrogram of topological overlap with colored bars to indicate module assignments. Dynamic tree cut shows the unmerged module assignments related to the merged dynamic for the final module colors.

**Supplementary Figure 4.**
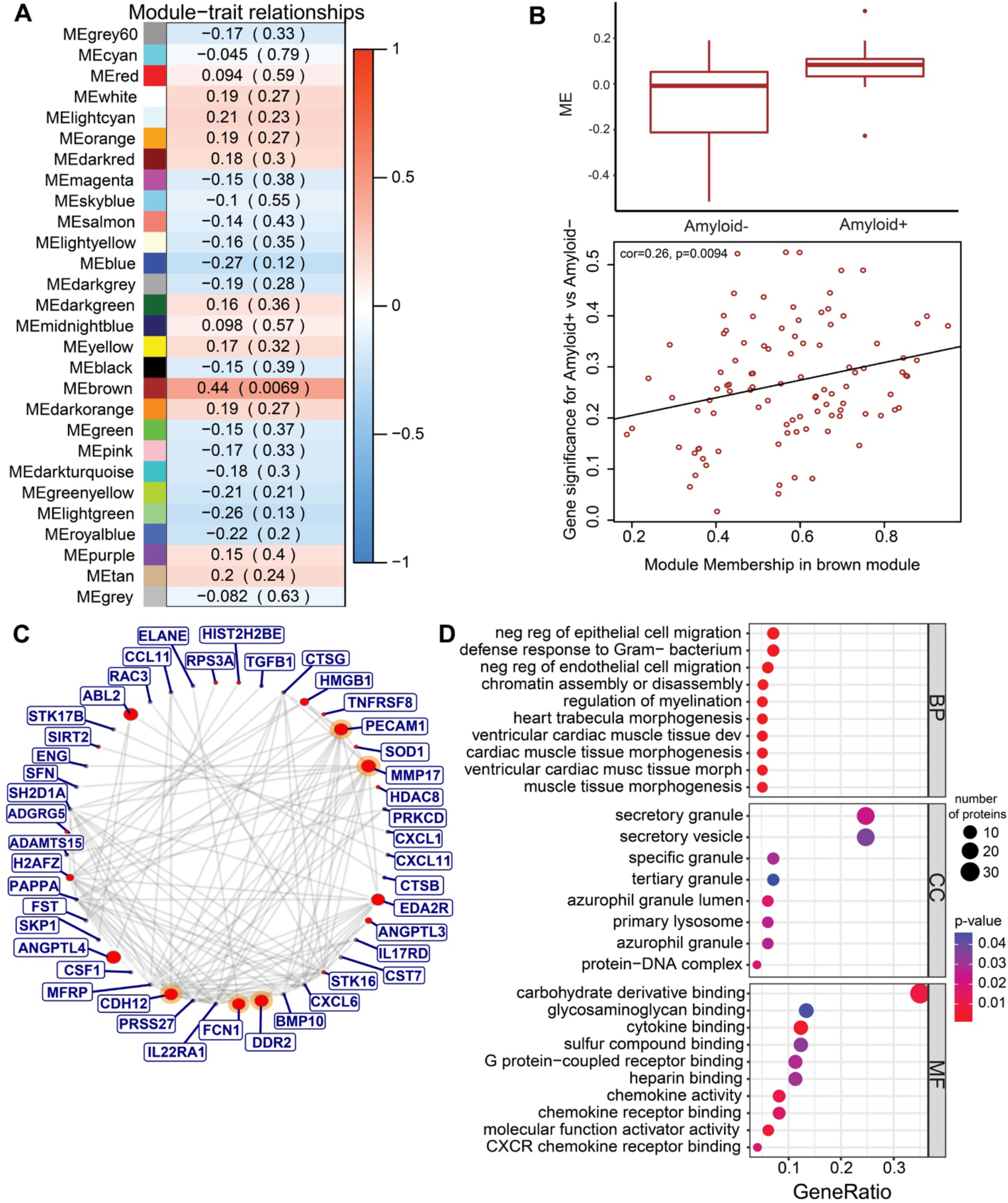
WGCNA module selection for co-regulatory changes between cognitively normal amyloid-positive and amyloid-negative subjects. **A**. Construction of the module-trait relationships plot for all module assignments (color labels) in addition to the module membership values and the respective module significance in parentheses. Selection of modules for further analysis is based on significance (*P* < 0.05). **B**. Bar graph demonstrating module eigengene differences for the brown module between amyloid-positive and amyloid-negative subjects. Scatterplot indicates the correlation among individual module eigen values and amyloid-positivity. **C**. Extraction of proteins from the brown module were visualized for protein-protein interactions by node and edge connectivity. Identification of hub genes (orange) was done by assessing the module membership values, while significantly upregulated proteins identified previously in Dream are highlighted in red. **D**. Enrichment analyses were performed using upregulated (amyloid-positive) proteins for Gene Ontology biological processes (BP), cellular components (CC), and molecular functions (MF).

